# When splicing is not all or none: GT>GC 5′ splice-site variants as a model for intermediate effects and challenges in variant classification

**DOI:** 10.1101/2025.10.30.685113

**Authors:** Jin-Huan Lin, Hao Wu, Xin-Ying Tang, Emmanuelle Masson, David N. Cooper, Claude Férec, Zhuan Liao, Wen-Bin Zou, Jian-Min Chen

**Affiliations:** Department of Gastroenterology, Changhai Hospital, Naval Medical University, Shanghai 200433, China; Shanghai Institute of Pancreatic Diseases, Shanghai 200433, China; Department of Gastroenterology, Eastern Hepatobiliary Surgery Hospital, Naval Medical University, Shanghai 200433, China; Univ Brest, Inserm, EFS, UMR 1078, GGB, 29200 Brest, France; Service de Génétique Médicale et de Biologie de la Reproduction, CHU Brest, 29200 Brest, France; Institute of Medical Genetics, School of Medicine, Cardiff University, Cardiff CF14 4XN, UK

**Keywords:** ACMG/AMP variant classification guidelines, ClinVar, Incomplete penetrance, Intermediate functional effects, Loss-of-function (LoF) variants, Missing heritability, Residual wild-type transcript, SpliceAI, Variant effect spectrum, Variants of uncertain significance (VUS)

## Abstract

Variants with intermediate functional effects—those that neither eliminate nor fully maintain normal gene function—represent an underrecognized source of genetic complexity and constitute a grey zone that challenges variant classification. Here, we address this issue using GT>GC (+2T>C) 5′ splice-site variants as a tractable model, as approximately 15% of such substitutions retain variable amounts of wild-type (WT) transcript. Using residual WT transcript as a quantitative readout, we first show that disease-associated GT>GC variants known to generate substantial WT transcript (for example, *SPINK1* c.194+2T>C, *HBB* c.315+2T>C, and *BRCA2* c.8331+2T>C) consistently complicate classification efforts. We then carried out a locus-wide assessment of all 26 GT>GC substitutions in *CFTR*, applying SpliceAI delta donor-loss scores as an initial screen and cross-checking classifications in expert-curated database. This evaluation indicated that three variants with low or intermediate SpliceAI scores are likely to retain WT transcript. Minigene analyses of four selected *CFTR* variants, together with additional examination of a *BAP1* GT>GC variant with conflicting clinical interpretations, illustrate the methodological challenges in predicting and experimentally quantifying residual WT transcript. These results emphasize that (i) *in silico* predictors cannot reliably discriminate between complete and partial splice disruption and (ii) experimental systems differ in their ability to detect subtle splicing outcomes. Overall, our integrative analysis shows that GT>GC variants generating appreciable WT transcript exemplify a broader class of intermediate-effect alleles that complicate both computational prediction and experimental assessment, underscoring the need for classification frameworks that incorporate quantitative functional data and better capture the continuum of variant effects.

## Introduction

Accurate classification of genetic variants remains a central challenge in human genetics.^1,2^ The increasing use of a genotype-first or genomic-first approach,^3,4^ enabled by exome and genome sequencing, has greatly magnified this challenge. In many cases, the mode of inheritance becomes apparent only after genetic analysis, and disorders presumed to follow a simple Mendelian pattern may instead have a complex, multifactorial, or oligogenic basis. This complexity underscores the limitations of the widely used American College of Medical Genetics and Genomics and the Association for Molecular Pathology (ACMG/AMP) framework.^5^ Its dichotomization of variants as either “pathogenic” or “benign,” although useful for clinical consistency, cannot account for the full spectrum of variant effects that contribute to human genetic disease.^3,6–9^ It should be noted that the current ACMG/AMP category “variant of uncertain significance (VUS)”^5^ is not a biological entity but a temporary designation expected to resolve as either pathogenic or benign once sufficient evidence becomes available.

For clarity, we focus here on variants that act through a loss-of-function (LoF) mechanism. In contrast to variants at the two extremes—those causing complete or near-complete loss of function of the affected allele, typically responsible for classic Mendelian disease phenotypes, and those retaining complete or near-complete normal function—variants with intermediate effects are challenging to interpret because they lie outside the scope of binary pathogenicity models. This challenge is illustrated by the multiple partial LoF variants identified in *SPINK1* (MIM: 167790).^8^ Despite their clear biological reality and clinical relevance, such variants remain underexplored and are not adequately addressed by any widely established classification frameworks, which offer limited guidance for interpreting quantitative or intermediate functional effects and thereby risk systematic over- or under-classification.

In this study, we explore GT>GC (+2T>C) 5′ splice-site variants as a tractable model to investigate how intermediate functional effects challenge classification. Although substitutions at canonical GT donor or AG acceptor sites typically abolish normal splicing,^10,11^ a subset of GT>GC substitutions defy this expectation. Specifically, a meta-analysis of 45 disease-associated variants, together with full-length gene splicing assay (FLGSA) data from 103 substitutions, showed that 15–18% retain residual wild-type (WT) transcript, in some cases at levels approaching 80% of normal.^12^ This variability can be clinically significant. For instance, in cystic fibrosis, as little as 5% normal cystic fibrosis transmembrane conductance regulator (CFTR) activity may be sufficient to prevent pulmonary disease,^13,14^ whereas in the hemophilias, factor VIII or IX levels above 5% may lead to only mild bleeding diatheses.^15,16^ Conversely, in disorders typically caused by complete LoF alleles, GT>GC variants may not be invariably pathogenic if substantial WT transcript is preserved, as we previously proposed.^12^

Using residual (“leaky”) WT transcript levels as a quantitative measure, this study illustrates the difficulties inherent in classifying GT>GC variants that generate substantial yet incomplete amounts of WT transcript. Our analysis highlights the challenges posed by intermediate functional effects and reinforces the need for classification frameworks capable of capturing the full continuum of variant effects.^6,8^

## Methods

### Identification and selection of GT>GC variants

Our 2019 study^12^ served as the starting point. From the FLGSA-analyzed GT>GC variants reported therein, we included all variants that both retained WT transcript and were registered in ClinVar^17^ (https://www.ncbi.nlm.nih.gov/clinvar/; accessed September 5, 2025) for analysis. To identify additional disease-associated GT>GC variants with evidence of residual WT transcript generation published since 2019, we conducted a Google-based literature search using the keyword “+2T>C,” complemented by cascade reference tracking and cross-referencing within the peer-reviewed literature (up to September 5, 2025). As in our previous work,^12^ we focused on studies in which RNA analysis was performed on material derived from affected individuals, and simple heterozygotes were excluded unless residual WT transcript generation from the variant allele was explicitly demonstrated by allele-specific reverse transcription-PCR (RT-PCR) in the original study. At the single-gene level, all 26 possible *CFTR* (MIM: 602421) GT>GC variants were subjected to initial evaluation. Finally, a *BAP1* (MIM: 603089) variant, c.783+2T>C (GenBank: NM_004656.4, NG_031859.1), identified during the literature search as having conflicting classifications,^18,19^ was included for additional minigene splicing analysis.

### *In silico* splicing predictions

*In silico* predictions were obtained using SpliceAI^20^ via the SpliceAI Lookup tool (https://spliceailookup.broadinstitute.org/; accessed September 5, 2025) with the following settings: genome version, hg38; GENCODE, basic; maximum distance, 10,000 bp; and REF & ALT scores displayed. The resulting delta donor loss (ΔDL) scores, derived from comparisons between WT and variant sequences, were used for analysis.

### Variant classification or description in public databases

Selected variants were queried in ClinVar (https://www.ncbi.nlm.nih.gov/clinvar/)^17^ to determine their current classification status. The Clinical and Functional TRanslation of CFTR database (CFTR2; http://www.cftr2.org/)^21^ and the CFTR-France database (https://cftr.chu-montpellier.fr/)^22^ were consulted to evaluate *CFTR* variant classifications. Variant information from the Cystic Fibrosis Mutation Database (CFTR1; http://www.genet.sickkids.on.ca/) was used to assist in the classification of two *CFTR* variants that were not registered in either CFTR2 or CFTR-France. Our evaluation of the variant classification or description data in these databases was last updated on October 28, 2025.

### Minigene splicing analysis of selected *CFTR* GT>GC variants

#### Variant selection

Of the 26 possible *CFTR* GT>GC variants (GenBank: NM_000492.4, NG_016465.4), we selected all three with low SpliceAI DL scores (≤0.56)—c.2490+2T>C, c.3717+2T>C and c.4242+2T>C—together with one variant (c.3873+2T>C) that had a high SpliceAI score (0.98) but was previously reported to generate WT transcript,^23^ for minigene splicing analysis.

#### Construction of pET01 and pSPL3 minigene expression vectors

For each selected +2T>C variant, the corresponding WT genomic sequences cloned into the pET01 and pSPL3 exon trapping vectors^24^ were always identical. Each *CFTR* WT insert comprised 213–217 bp from the 3′ end of intron N−1, the entire exon N, and 213–216 bp from the 5′ end of intron N (N = variant-affected intron). Cloning of WT constructs and introduction of the selected +2T>C variants, followed by Sanger confirmation, were performed by GENEWIZ Biotech Co. (Suzhou, China).

#### Cell culture, transfection, RT-PCR, and sequencing

Experiments were carried out essentially as previously described.^24^ Human embryonic kidney 293T (HEK293T) cells were maintained in DMEM (Gibco) supplemented with 10% fetal calf serum (Procell). Cells (3.5 × 10⁵/well) were seeded in 6-well plates 24 h before transfection. Per well, 2.5 µg WT or variant plasmid were transfected using HieffTrans Universal Transfection Reagent (Yeasen). Forty-eight hours later, total RNA was extracted with the FastPure Cell/Tissue Total RNA Isolation Kit V2 (+gDNA wiper) (Vazyme). RT was carried out using the HiScript III 1st Strand cDNA Synthesis Kit (Vazyme), incorporating 2 µL 5×gDNA wiper Mix, 2 µL 10×RT Mix, 2 µL HiScript III Enzyme Mix, 1 µL Oligo (dT)20VN, and 1 µg total RNA. RT-PCR was performed in a 25-μL reaction mixture containing 12.5 μL 2×Taq Master Mix (Vazyme), 1 μL cDNA, and 0.4 μM each primer. The primer pairs were as follows: pET01 vector—forward 5′-GAGGGATCCGCTTCCTGGCCC-3′, reverse 5′-CTCCCGGGCCACCTCCAGTGCC-3′; pSPL3 vector—forward 5′-TCTGAGTCACCTGGACAACC-3′, reverse 5′-ATCTCAGTGGTATTTGTGAGC-3′. The PCR program comprised an initial denaturation step at 94°C for 5 min, followed by 35 cycles of denaturation at 94°C for 30 s, annealing at 55°C for 30 s, and extension at 72°C for 1 min, and a final extension step at 72°C for 7 min. RT-PCR products were separated on agarose gels, excised, and purified (Omega Bio-Tek Gel Extraction Kit). Sequencing primers were the same as those used for RT-PCR, and reactions were performed with the BigDye Terminator v3.1 Cycle Sequencing Kit (Applied Biosystems).

### Full-length and minigene splicing analysis of *BAP1* c.783+2T>C

The full-length genomic *BAP1* WT insert, spanning from the translation initiation codon to the termination codon (GenBank: NM_004656.4, NG_031859.1), was amplified from genomic DNA of a normal control using long-range PCR. The resulting fragment was cloned into the pcDNA3.1/V5-His-TOPO vector. These procedures, as well as subsequent transfection and RT–PCR, were performed essentially as previously described.^12^

Minigene splicing analysis followed the same procedures as for *CFTR* variants, except that two WT constructs (and their corresponding variant constructs) were generated. The *BAP1* Exon 9 insert comprised 337 bp from intron 8, exon 9, and 302 bp from intron 9, whereas the *BAP1* Exon 8–10 insert comprised 197 bp from intron 7, exon 8, intron 8, exon 9, intron 9, exon 10, and 243 bp from intron 10.

## Results

### Analytical approach

Building on our 2019 findings that some GT>GC variants retain WT transcript,^12^ we analyzed three successive variant categories (Figure 1). First, we re-examined variants previously characterized by FLGSA. Second, we considered disease-associated GT>GC variants reported since 2019 that generate WT transcript. Third, we evaluated all possible GT>GC variants in *CFTR* and functionally tested four selected variants. Across these categories, we assessed residual WT transcript levels, genetic and clinical findings, SpliceAI ΔDL scores, and database classifications, where available. In addition, we conducted minigene splicing analysis of the *BAP1* c.783+2T>C variant, which has conflicting classifications.^18,19^ Integrating these findings, we delineated the classification challenges posed by variants with intermediate functional effects—hereafter referred to conceptually as grey-zone variants.

**Figure 1.**
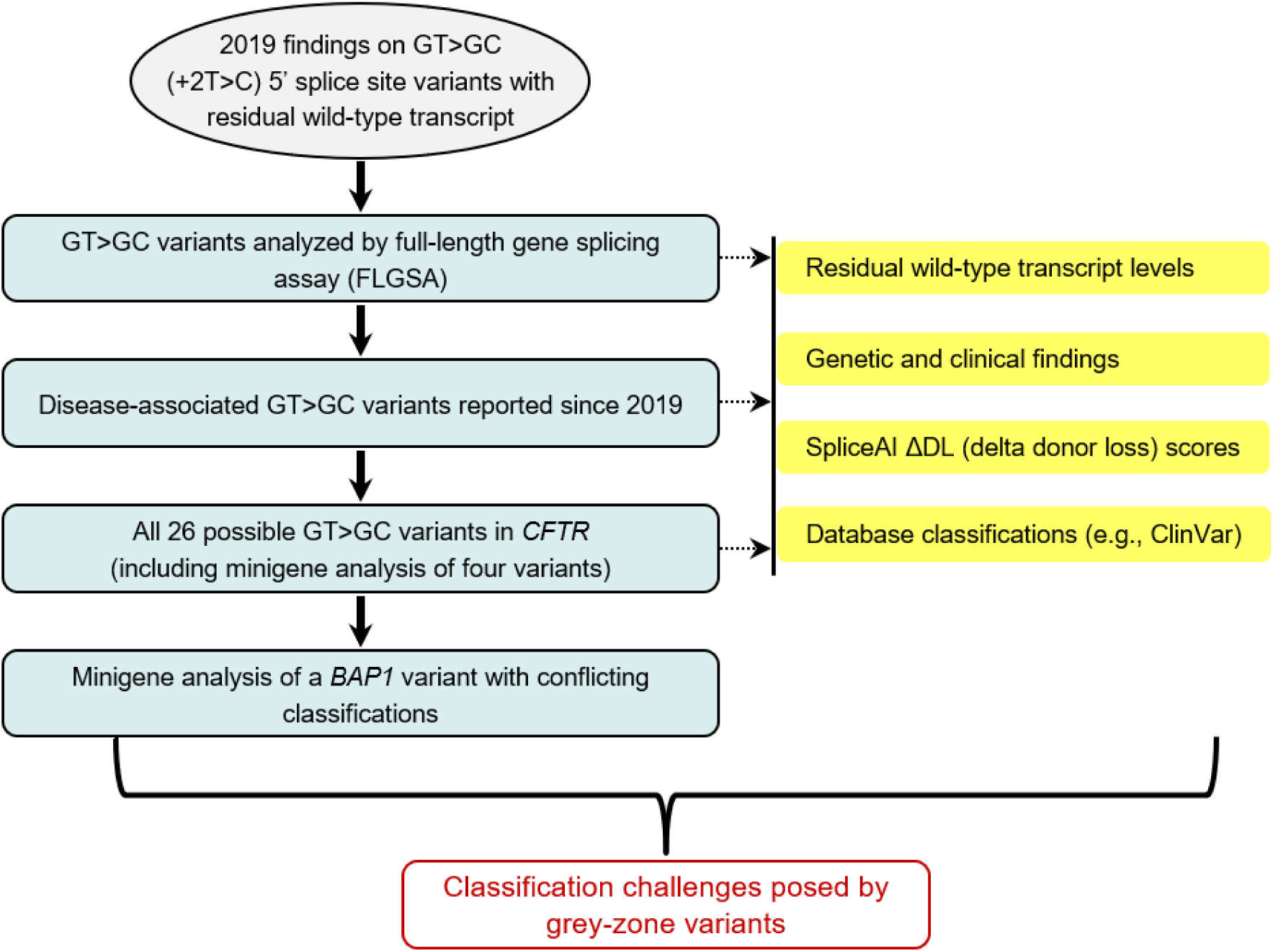
Scheme of the analytical approach. GT>GC variants were analyzed in three successive categories, together with minigene analysis of a *BAP1* variant with conflicting classifications. Within each category, variants were examined for four features (yellow shading). The red box highlights the central issue addressed in this study—the classification challenges posed by grey-zone variants that retain substantial yet incomplete wild-type transcript generation.

### Comparing disease-associated and FLGSA-analyzed GT>GC variants

In our previous study,^12^ the proportion of GT>GC variants retaining WT transcript was similar between disease-associated and FLGSA-analyzed variants: 15.6% (7 of 45) versus 18.4% (19 of 103). The magnitude of residual WT transcript levels, however, differed markedly. The seven disease-associated variants yielded only low levels, from trace amounts up to 15% normal, whereas of the ten variants generating exclusively WT transcript (as judged by gel electrophoresis), eight retained levels ranging from 13.7% to 84% of normal, with only two falling below 5%.

The consistently low WT transcript levels observed for the seven disease-associated variants most likely reflect clinical ascertainment bias, as noted previously.^12^ Nonetheless, of the four variants retaining 10–15% WT transcript, three—*CAV3* (MIM: 601253) c.114+2T>C (GenBank: NM_033337.3, NG_008797.2),^25^ *CD40LG* (MIM: 300386) c.346+2T>C (GenBank: NM_000074.3, NG_007280.1),^26^ and *DMD* (MIM: 300377) c.8027+2T>C (GenBank: NM_004006.3, [NG_012232.1](https://www.ncbi.nlm.nih.gov/nuccore/NM_004006.3))^27^—were associated with milder clinical phenotypes; the remaining variant, *SPINK1* c.194+2T>C (GenBank: NM_001379610.1, NG_008356.2), is discussed below.

The corresponding WT constructs for the 103 FLGSA-analyzed substitutions were invariably preselected to yield a single or quasi-single RT–PCR band of the expected WT transcript size, consistent with their respective reference mRNA sequences.^12^ We also analyzed *HBB* (MIM: 141900) c.315+2T>C (GenBank: NM_000518.5, NG_059281.1),^28^ despite its WT construct producing two strong bands—one corresponding to the WT transcript and the other representing an aberrantly spliced transcript. This variant also generated WT transcript in FLGSA,^12^ bringing the total number of WT transcript–generating GT>GC variants identified by FLGSA to 20.

### Detailed analysis of three FLGSA-analyzed GT>GC variants

Of the 20 GT>GC variants previously shown by FLGSA^12^ to generate residual WT transcript, only three were found to be registered in ClinVar^17^ (https://www.ncbi.nlm.nih.gov/clinvar/; assessed September 5, 2025). These three variants, described below, illustrate how the presence of residual WT transcript can challenge variant classification.

#### SPINK1 c.194+2T>C

Among GT>GC variants, *SPINK1* c.194+2T>C (GenBank: NM_001379610.1, NG_008356.2) is distinctive in that transcript expression has been assessed both *in vivo* (using RNA from an affected homozygote)^29^ and *in vitro* using FLGSA.^12^ Both approaches yielded concordant results, with approximately 10% of normal WT transcript levels from the variant allele.^30^ SpliceAI predicts a ΔDL score of 0.33, consistent with a substantial yet incomplete reduction in normal splicing.

Our in-depth analysis of *SPINK1*-related chronic pancreatitis^8^ indicated that this ∼10% residual WT transcript places c.194+2T>C in the grey zone, conferring a well-defined genetic effect that is too weak for “pathogenic” under current ACMG/AMP criteria^5^ but too strong for the proposed “risk allele” category.^7^ We therefore suggested classifying it as “predisposing,” reflecting its partial LoF nature and intermediate genetic impact.^8^ ClinVar lists this variant as pathogenic/likely pathogenic ([VCV000132142.62](https://www.ncbi.nlm.nih.gov/clinvar/variation/132142/); accessed October 28, 2025), but convergent evidence from functional assays, *in silico* prediction, and clinical and genetic data supports its role as proof of concept highlighting the need for expanded classification frameworks.^8^

#### HBB c.315+2T>C

This variant was identified during routine thalassemia screening in a 34-year-old individual who exhibited hypochromia and microcytosis consistent with a thalassemia trait rather than β-thalassemia major. The individual was compound heterozygous for *HBB* c.315+2T>C and *HBB* c.79G>A (p.Glu27Lys) (GenBank: NM_000518.5, NG_059281.1).^28^ The original authors reported that *HBB* c.315+2T>C had a limited effect on splicing efficiency, stating that it “results in expression that is approximately two-thirds of the normal allele.”^28^ However, this estimate was based on hemoglobin quantitation and was likely confounded by the *trans*-inherited *HBB* c.79G>A (p.Glu27Lys) variant. The latter likely retains substantial normal function, as most individuals homozygous for *HBB* c.79G>A (p.Glu27Lys) exhibit only mild hypochromic microcytic anemia.^31^

Our FLGSA analysis confirmed that *HBB* c.315+2T>C allows appreciable production of WT transcript.^12^ Here, using ImageJ (https://imagej.net/ij/), we compared the intensity of the WT transcript band derived from the variant allele with that from the WT allele in the original gel.^12^ The variant allele was estimated to retain approximately 30% of WT transcript output, providing a more direct and reliable measure of its residual transcript level. Notably, the orthologous rabbit variant also generated WT transcripts,^32,33^ supporting conservation of the partial splicing effect across species. The SpliceAI ΔDL score for *HBB* c.315+2T>C was 0.72.

*HBB* alleles in β-thalassemia have traditionally been classified as β⁰ (complete loss of β-globin synthesis), β⁺ (moderate-to-severe reduction), or β⁺⁺ (“silent,” minimal reduction).^34^ *HBB* c.315+2T>C was originally described as a mild β⁺ allele,^28^ consistent with our FLGSA finding of significant but not complete loss of WT transcript.^12^ Nevertheless, it is currently annotated in ClinVar as “pathogenic” ([VCV000869355.2](https://www.ncbi.nlm.nih.gov/clinvar/variation/869355/); accessed October 28, 2025)—a designation that may warrant re-evaluation in view of its substantial residual WT transcript generation and the mild hematological phenotype observed in the 34-year-old compound heterozygote.

#### MGP c.94+2T>C

*MGP* (MIM: 154870) c.94+2T>C (GenBank: NM_000900.5, NG_023331.2) was shown by FLGSA^12^ to retain approximately 80% of WT transcript, whereas SpliceAI predicted a strong splice-disrupting effect (ΔDL score, 0.97). Despite this discrepancy, ClinVar lists *MGP* c.94+2T>C as “likely pathogenic” ([VCV001987659.2](https://www.ncbi.nlm.nih.gov/clinvar/variation/1987659/); accessed October 28, 2025), based on a single clinical testing submission (SCV003031993.2) with unknown affected status and no published case reports. This classification appears to rest on four implicit assumptions:

1. the variant alters a canonical donor splice site;
2. such variants generally disrupt splicing;
3. disruption of canonical sites typically abolishes protein function; and
4. LoF variants in *MGP* are known to be pathogenic.

As this example shows, ClinVar classifications may at times rely on inferred reasoning rather than direct experimental evidence.

### Disease-associated GT>GC variants reported since 2019

We next examined disease-associated GT>GC variants reported since 2019, focusing on those with WT transcript generation documented in samples derived from affected individuals. Among the five variants identified, *SBDS* (MIM: 607444) c.258+2T>C (GenBank: NM_016038.4, NG_007277.1)^35^ and *SRP68* (MIM: 604858) c.184+2T>C (GenBank: NM_014230.4, NC_000017.11),^36^ produced only trace (∼2%) or low amounts of WT transcript, with SpliceAI ΔDL scores of 0.93 and 0.37, respectively. *TNXB* (MIM: 600985) c.12469+2T>C (GenBank: NM_001365276.2, NG_008337.2) retained a moderate level of WT transcript by allele-specific RT–PCR,^37^ consistent with its very low ΔDL score of 0.01, and has been associated with moderate Ehlers–Danlos–like manifestations in congenital adrenal hyperplasia.^37^ The remaining two variants—*BRCA2* (MIM: 600185) c.8331+2T>C (GenBank: NM_000059.4, [NG_012772.3](https://www.ncbi.nlm.nih.gov/nuccore/NM_000059.4)) and *DOCK8* (MIM: 611432) c.3234+2T>C (GenBank: NM_203447.4, NG_017007.1)—are discussed below.

#### BRCA2 c.8331+2T>C

*BRCA2* c.8331+2T>C (GenBank: NM_000059.4, NG_012772.3) is associated with a high SpliceAI ΔDL score (0.93) and is listed in ClinVar as “pathogenic/likely pathogenic” ([VCV000267692.71](https://www.ncbi.nlm.nih.gov/clinvar/variation/267692/); accessed October 28, 2025). However, allele-specific RT–PCR in a heterozygous carrier demonstrated approximately 60% of WT transcript from the variant allele.^38^ A validated *BRCA1/2* history-weighting algorithm, which compares cancer histories with expectations for pathogenic versus benign variants,^39^ yielded a benign call.^38^ Moreover, *BRCA2* appears relatively tolerant of reduced transcript dosage, and one study suggested that variants producing more than 5% WT transcript should, in the absence of other supporting evidence, be considered VUS.^40^ While the original report reclassified c.8331+2T>C from “pathogenic” to a “VUS,”^38^ the combined evidence appears to point toward a benign interpretation under current ACMG/AMP principles.

#### *DOCK8* c.3234+2T>C

Biallelic LoF variants in *DOCK8* cause autosomal recessive hyper-IgE syndrome-2 (MIM: 243700). *DOCK8* c.3234+2T>C (GenBank: NM_203447.4, NG_017007.1) was reported in the homozygous state in an infant with severe viral infections.^41^ T cells from this individual showed a predominance of WT transcript, normal DOCK8 protein accumulation, intact cellular function, and no DOCK8 deficiency phenotype. The clinical presentation was instead explained by a homozygous *IFNAR1* (MIM: 107450) nonsense variant (c.1156G>T, p.Glu386Ter; GenBank: NM_000629.3).^41^ Despite this evidence for near-complete WT transcript generation, lack of disease relevance, and a modest SpliceAI ΔDL score (0.49), three of the four ClinVar submissions for *DOCK8* c.3234+2T>C ([VCV000573664.15](https://www.ncbi.nlm.nih.gov/clinvar/variation/573664/); accessed October 28, 2025) classified it as “likely pathogenic,” while one classified it as “benign.”

*BRCA2* c.8331+2T>C and *DOCK8* c.3234+2T>C exemplify how variants predicted—or assumed—to be fully disruptive can nevertheless yield intermediate or even near-complete levels of WT transcript, underscoring the need for direct transcript-level evidence for accurate classification. These two variants are summarized in Table 1 alongside the three FLGSA-analyzed examples (*SPINK1* c.194+2T>C, *HBB* c.315+2T>C, and *MGP* c.94+2T>C).

**Table 1.** Five GT>GC variants with residual WT transcript that present classification challenges.

| Variant<sup>a</sup> | Reference mRNA and genomic sequences | hg38 coordinate | Residual WT transcript<sup>b</sup> | Assessment method | SpliceAI ΔDL score | Clinical/genetic findings | ClinVar classification |
| --- | --- | --- | --- | --- | --- | --- | --- |
| *SPINK1*<br>c.194+2T>C | NM_001379610.1,<br>NG_008356.2 | chr5-147828020-A-G | 10% | RT-PCR of material from an affected homozygote;<sup>29</sup> FLGSA<sup>12</sup> | 0.33 | Well-defined genetic effect, too weak for pathogenic classification<sup>8</sup> | Pathogenic/likely pathogenic |
| *HBB*<br>c.315+2T>C | NM_000518.5,<br>NG_059281.1 | chr11-5226575-A-G | 30% | FLGSA<sup>12</sup> | 0.72 | Originally described as a mild $\beta^+$-thalassemia allele<sup>28</sup> | Pathogenic |
| *MGP*<br>c.94+2T>C | NM_000900.5,<br>NG_023331.2 | chr12-14884211-A-G | 80% | FLGSA<sup>12</sup> | 0.97 | Not reported in affected individuals | Likely pathogenic |
| *BRCA2*<br>c.8331+2T>C | NM_000059.4,<br>NG_012772.3 | chr13-32363535-T-C | 60% | Allele-specific RT-PCR of affected individual-derived material<sup>38</sup> | 0.93 | *BRCA2* history-weighting algorithm<sup>39</sup> indicated a benign profile;<sup>38</sup> *BRCA2* exhibits tolerance to reduced transcript dosage<sup>40</sup> | Pathogenic/likely pathogenic |
| *DOCK8*<br>c.3234+2T>C | NM_203447.4,<br>NG_017007.1 | chr9-399261-T-C | Predominant WT | RT-PCR and protein analyses of T cells from an affected homozygote<sup>41</sup> | 0.49 | Disease explained by an unlinked homozygous *IFNAR1* nonsense variant<sup>41</sup> | Conflicting classifications of pathogenicity |

Abbreviations: ΔDL, delta donor loss; FLGSA, full-length gene splicing assay; RT-PCR, reverse transcription-PCR; WT, wild-type.
<sup>a</sup>Variants are arranged in order of mention in the main text.
<sup>b</sup>Percentage of WT transcript generated from the variant allele relative to that from the WT allele.

### Locus-wide analysis of *CFTR* GT>GC variants

Having examined GT>GC variants that retain WT transcript generation across multiple genes, we next asked how frequent and systematic this phenomenon might be within a single locus. We selected *CFTR* (GenBank: NM_000492.4, NG_016465.4), in which biallelic LoF variants cause cystic fibrosis (CF; MIM: 219700),^42,43^ as a model system because it harbors 26 GT donor splice sites (Table S1), is among the most extensively studied disease-related genes, and benefits from expert-curated databases CFTR2^21^ and CFTR-France^22^ that provide authoritative variant annotations.

Building upon our previous^12,30^ and current findings, we observed that although some GT>GC variants retaining WT transcript are associated with high SpliceAI ΔDL scores, most have low scores. We therefore used SpliceAI ΔDL scores as the starting point for this analysis, complemented by cross-examination of variant classifications in CFTR2 (http://www.cftr2.org/; accessed October 28, 2025) and CFTR-France (https://cftr.chu-montpellier.fr/; accessed October 28, 2025) (Figure 2A). Notably, all 16 *CFTR* GT>GC variants that we grouped as “pathogenic” had SpliceAI ΔDL scores greater than 0.80. These included 13 designated as “CF-causing” in CFTR2, one absent from CFTR2 but listed as “likely pathogenic” in CFTR-France (c.3468+2T>C), and two absent from both CFTR2 and CFTR-France but classified by us as “pathogenic” based on clinical and genetic data in CFTR1 (http://www.genet.sickkids.on.ca/; accessed October 28, 2025) (c.2908+2T>C and c.2988+2T>C). Eight *CFTR* GT>GC variants were not registered in these three databases, of which seven also scored above 0.80 and one had a score of 0.56 (c.3717+2T>C). The remaining two *CFTR* GT>GC variants—one listed as a VUS in CFTR-France (c.2490+2T>C) and one reported in CFTR1 in two individuals with suspected CF (c.4242+2T>C)—had SpliceAI ΔDL scores below 0.56 (Figure 2A,B). Table S1 provides full details of all 26 *CFTR* GT>GC variants, including the rationale for our classifying two as “pathogenic.”

**Figure 2.**
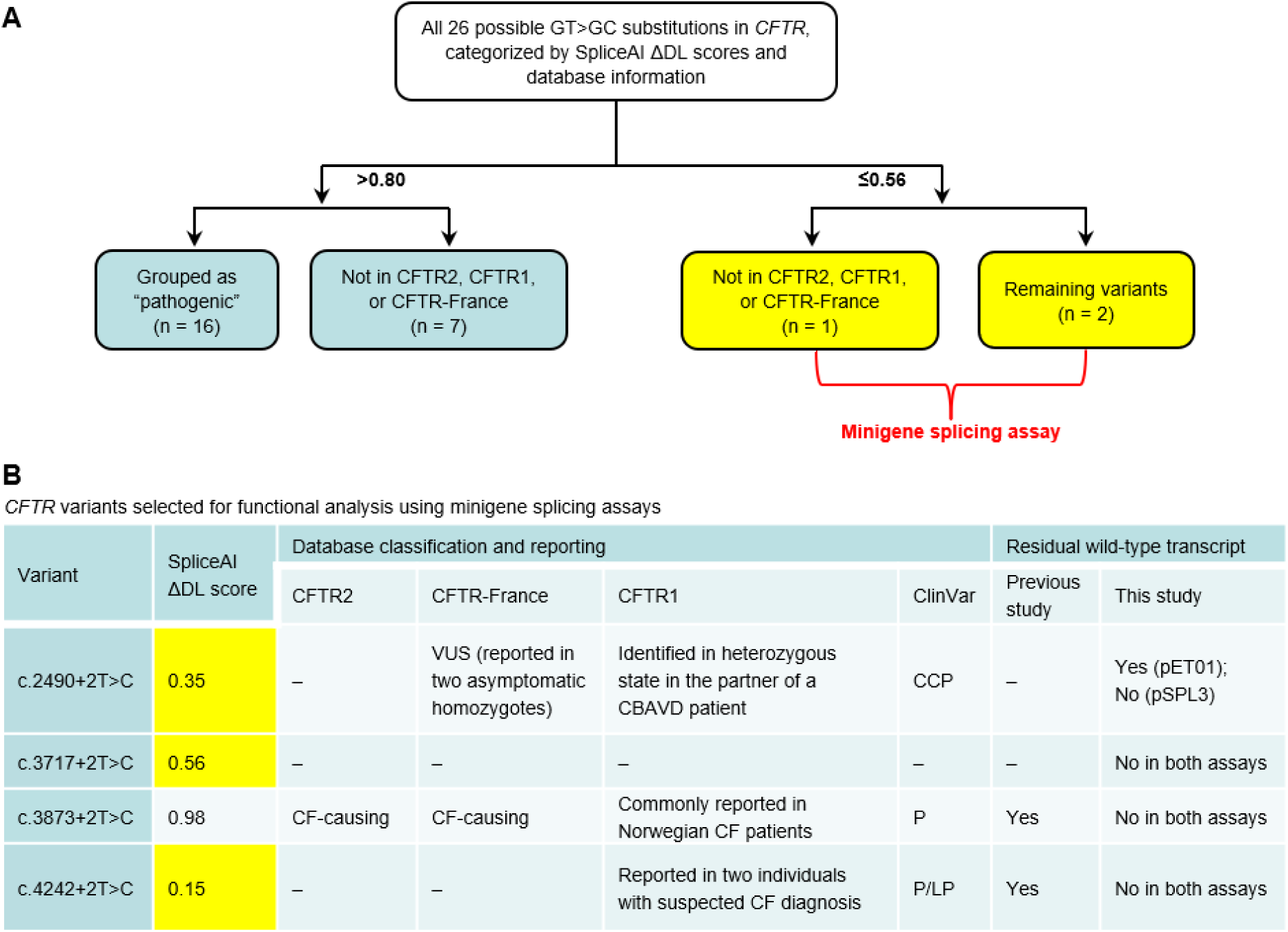
Analysis of GT>GC substitutions in *CFTR*. **(A)** Flowchart summarizing the 26 possible *CFTR* GT>GC substitutions categorized by SpliceAI ΔDL scores and database information (see Table S1 for details). Variants with low scores (≤0.56) were selected for minigene splicing assays. **(B)** Four variants analyzed by minigene splicing assays. In “Database classification and reporting,” note that, unlike CFTR2 and CFTR-France, CFTR1 does not classify registered variants. In “Residual wild-type transcript,” “Yes” and “No” indicate whether the variant allele generated wild-type transcript. Results from this study are shown separately for the two assay systems (pET01 and pSPL3; see Figure 3), with “No in both assays” indicating absence of wild-type transcript from the variant allele in both systems. “Previous study” refers to Joynt et al.^23^ Abbreviations: CBAVD, congenital bilateral absence of the vas deferens; CF, cystic fibrosis; CCP, conflicting classifications of pathogenicity; ΔDL, delta donor loss; P, pathogenic; P/LP, pathogenic/likely pathogenic; VUS, variant of uncertain significance.

Assuming that the three variants with relatively low SpliceAI ΔDL scores (c.2490+2T>C, 0.35; c.3717+2T>C, 0.56; and c.4242+2T>C, 0.15) generate WT transcript, at least 11.5% of the 26 *CFTR* GT>GC variants would retain WT transcript. This proportion is broadly consistent with the 15–18% frequency observed across genes in our previous study.^12^

Among these three *CFTR* variants with relatively low SpliceAI ΔDL scores, only c.4242+2T>C had previously undergone functional analysis. Using an expression minigene (EMG) system comprising the full *CFTR* coding sequence and intronic regions tailored to the variant under study, generation of residual WT transcript from the c.4242+2T>C allele was confirmed by Sanger sequencing, and mature CFTR protein was detected in transfected cells by immunoblotting.^23^ The only other *CFTR* GT>GC variant reported to yield WT transcripts is c.3873+2T>C, although this was mentioned only briefly and without accompanying experimental data.^23^ Notably, c.3873+2T>C is classified as “CF-causing” in both CFTR2 and CFTR-France, consistent with its high SpliceAI ΔDL score (0.98) (Figure 2B).

We tested the three variants with low SpliceAI ΔDL scores, together with c.3873+2T>C, in minigene assays using both pET01 and pSPL3 vectors (Figure 3; see also Figure S1). Findings for each variant are detailed below.

**Figure 3.**
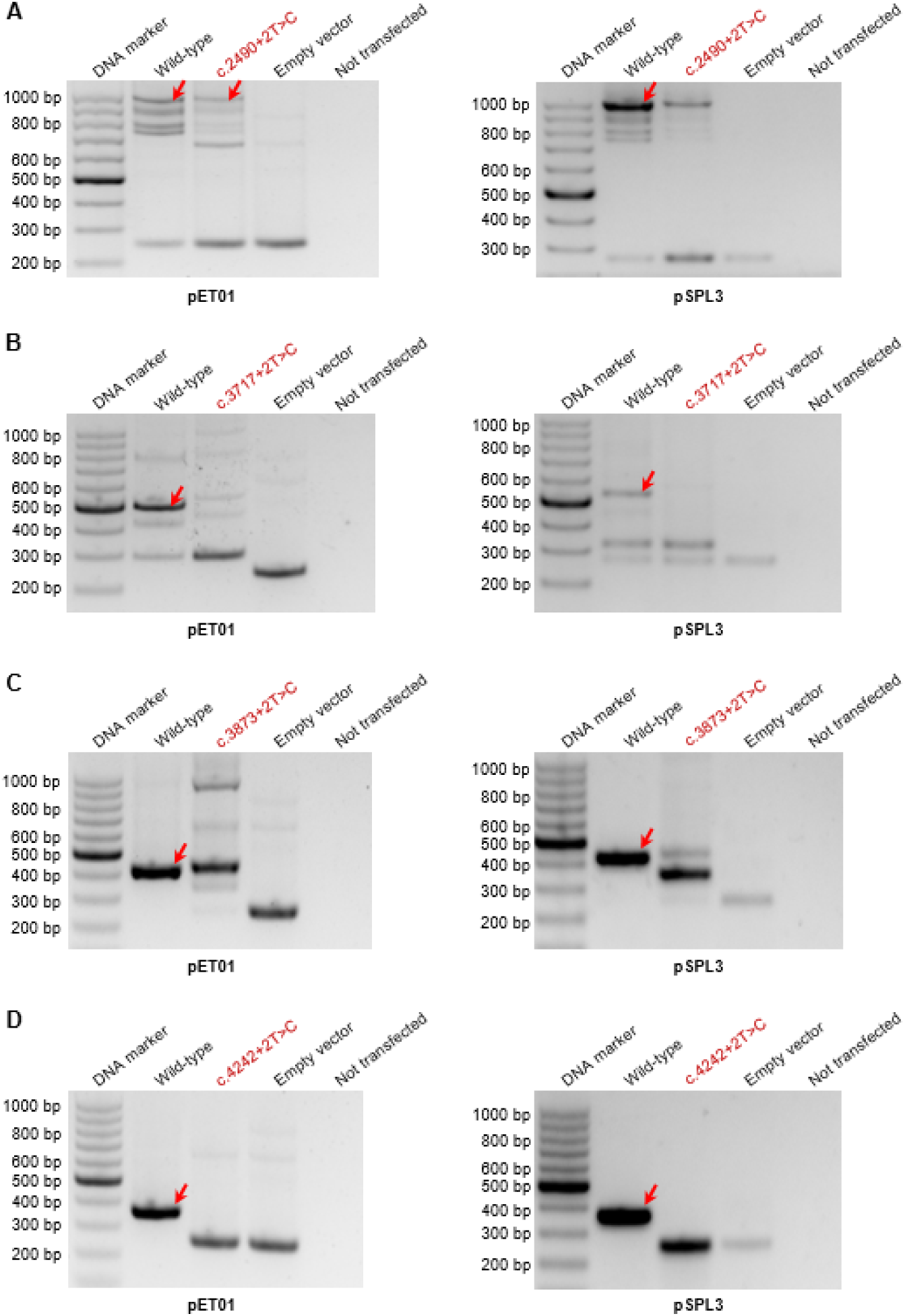
Minigene splicing analysis of four selected *CFTR* GT>GC substitutions using pET01 and pSPL3 vectors. Each variant was analyzed with its preceding exon and short flanking intronic sequences cloned into the respective reporter vectors. Arrows indicate bands corresponding to the normally spliced transcript, in accordance with the *CFTR* mRNA reference sequence NM_000492.4. Additional bands are annotated in Figure S1.

#### c.2490+2T>C (intron 14)

WT constructs in both minigene vectors generated multiple RT–PCR products, including the WT transcript. Importantly, WT transcript from the variant allele was detected in the pET01 vector (Figure 3A), consistent with its modest SpliceAI ΔDL score (0.35). Although this variant is not classified by CFTR2, it is listed in CFTR-France as a VUS, having been reported in two asymptomatic homozygotes. In CFTR1, it was described in the heterozygous state in the partner of an individual affected by congenital bilateral absence of the vas deferens (Figure 2B). Furthermore, ClinVar lists this variant under “conflicting classifications of pathogenicity” ([VCV000370078.23](https://www.ncbi.nlm.nih.gov/clinvar/variation/370078/); accessed October 28, 2025), based on six clinical-testing submissions (“pathogenic” × 3, “likely pathogenic” × 2, and “uncertain significance” × 1). Only one submission specified a known affected status—homozygosity for c.2490+2T>C, labeled “pathogenic,” in an individual with fetal cystic hygroma (SCV001737053.1). Another submission, designated “likely pathogenic” (SCV001363801.5), explicitly noted that “no occurrence of c.2490+2T>C in individuals affected with cystic fibrosis and no experimental evidence demonstrating its impact on protein function have been reported.” Taken together, the clinical, genetic, computational, and experimental evidence indicate that c.2490+2T>C is unlikely to represent a pathogenic *CFTR* variant in the context of classic CF.

#### c.3717+2T>C (intron 22)

No WT transcript from the variant allele was detected in either minigene vector (Figure 3B), despite its intermediate SpliceAI ΔDL score (0.56). This variant is not listed in CFTR2, CFTR-France, CFTR1, or ClinVar (Figure 2B).

#### c.3873+2T>C (intron 23)

No WT transcript from the variant allele was observed in either vector (Figure 3C); this finding is consistent with its high SpliceAI ΔDL score (0.98), its designation as “CF-causing” in both CFTR2 and CFTR-France, and its “pathogenic” classification in ClinVar (<u>VCV000053828.18</u>; accessed October 28, 2025) (Figure 2B). Moreover, consistent with its description in CFTR1, this variant is among the *CFTR* alleles recurrently identified in individuals with clinically diagnosed CF in Norway.^44^ Taken together, c.3873+2T>C represents a pathogenic *CFTR* variant causing classic CF. The level of WT transcript generated from the c.3873+2T>C allele, as previously reported,^23^ is presumably minimal.

#### c.4242+2T>C (intron 26)

No WT transcript from the variant allele was detected in either minigene vector (Figure 3D), in contrast to the earlier EMG report of residual WT transcript and protein production^23^ and to its very low SpliceAI ΔDL score (0.15). Notably, this variant is not listed in CFTR2 or CFTR-France but appears in CFTR1 as present in two individuals with suspected cystic fibrosis (Figure 2B). ClinVar lists c.4242+2T>C as “pathogenic/likely pathogenic” ([VCV000035884.8](https://www.ncbi.nlm.nih.gov/clinvar/variation/35884/); accessed October 28, 2025), based on three submissions (“pathogenic” × 1 and “likely pathogenic” × 2), all from clinical genetic testing laboratories. The collection method was described as “clinical testing” for two submissions (SCV002630543.2 and SCV004675881.2), both with unknown affected status, and as “curation” for the remaining submission (SCV000052191.3). Taken together, this variant may best be classified as a VUS at present.

In summary, locus-wide analysis of *CFTR* confirms that a subset of GT>GC variants retain WT transcript. Functional testing of selected *CFTR* GT>GC variants highlights both inter-assay variability and the inherent limitations of minigene systems in fully recapitulating native splicing.^24^

### Challenges of splicing assays: the case of *BAP1* c.783+2T>C

As illustrated by *CFTR* variants, *in vitro* splicing assays may fail to yield clear or consistent results, complicating variant classification. This challenge is further exemplified by *BAP1* c.783+2T>C (GenBank: NM_004656.4, NG_031859.1). Because heterozygous germline LoF variants in *BAP1* underlie tumor predisposition syndrome 1 (TPDS1; MIM: 614327) and *BAP1* c.783+2T>C affects the canonical intron 9 donor splice site, this variant was initially classified in ClinVar as “likely pathogenic,” as noted by Goldberg et al.^18^ These authors reclassified it as a VUS based on the following considerations:

1. the variant was identified in seven families (six Jewish Iraqi and one non-Jewish) with nonclassical *BAP1*-related TPDS1 tumor types;
2. the variant exhibited low penetrance;
3. the variant had a relatively high allele frequency in Jewish Iraqi individuals (3.6%);
4. the variant had a low SpliceAI score (0.08); and
5. no aberrantly spliced transcripts were detected by RT-PCR using RNA from blood cells of two probands.

A subsequent study^19^ identified a methodological limitation: the forward primer used by Goldberg et al.^18^ was located within exon 9, rendering it incapable of detecting exon 9 skipping. Using a primer in exon 8, Sculco et al.^19^ analyzed *BAP1* c.783+2T>C heterozygotes and observed an alternative splicing pattern consistent with exon 9 skipping. These authors proposed reinstating a “likely pathogenic” classification while acknowledging that the effect of *BAP1* c.783+2T>C on generating aberrant transcripts might be incomplete.

To further clarify this issue, we performed additional splicing assays (Figure 4; Figure S2). We first attempted FLGSA, as the *BAP1* genomic sequence from start to stop codon (<8 kb) falls within the upper size limit of the insert cloned into the pcDNA3.1/V5-His-TOPO vector routinely used in this assay.^12^ However, the WT construct, when transfected into HEK293T cells, failed to yield clear RT-PCR bands (Figure 4A). Notably, in our previous study, only 33 of 119 (28%) WT full-length constructs produced a single or quasi-single band of the expected size after transfection.^12^ We therefore turned to minigene assays using pET01 and pSPL3 vectors. WT constructs spanning genomic sequence from exon 8 to exon 10 (including exon 8, intron 8, exon 9, intron 9, and exon 10) with short flanking sequences from introns 7 and 10 produced multiple RT-PCR bands, none corresponding to the WT transcript defined by NM_004656.4; the corresponding variant constructs also failed to generate a normally spliced transcript, although the banding patterns differed from those of the WT (Figure 4B). In contrast, shorter constructs containing exon 9 with short flanking intronic sequences generated the expected WT band from the WT allele, whereas only transcripts lacking exon 9 were produced from the variant allele (Figure 4C; Figure S2).

**Figure 4.**
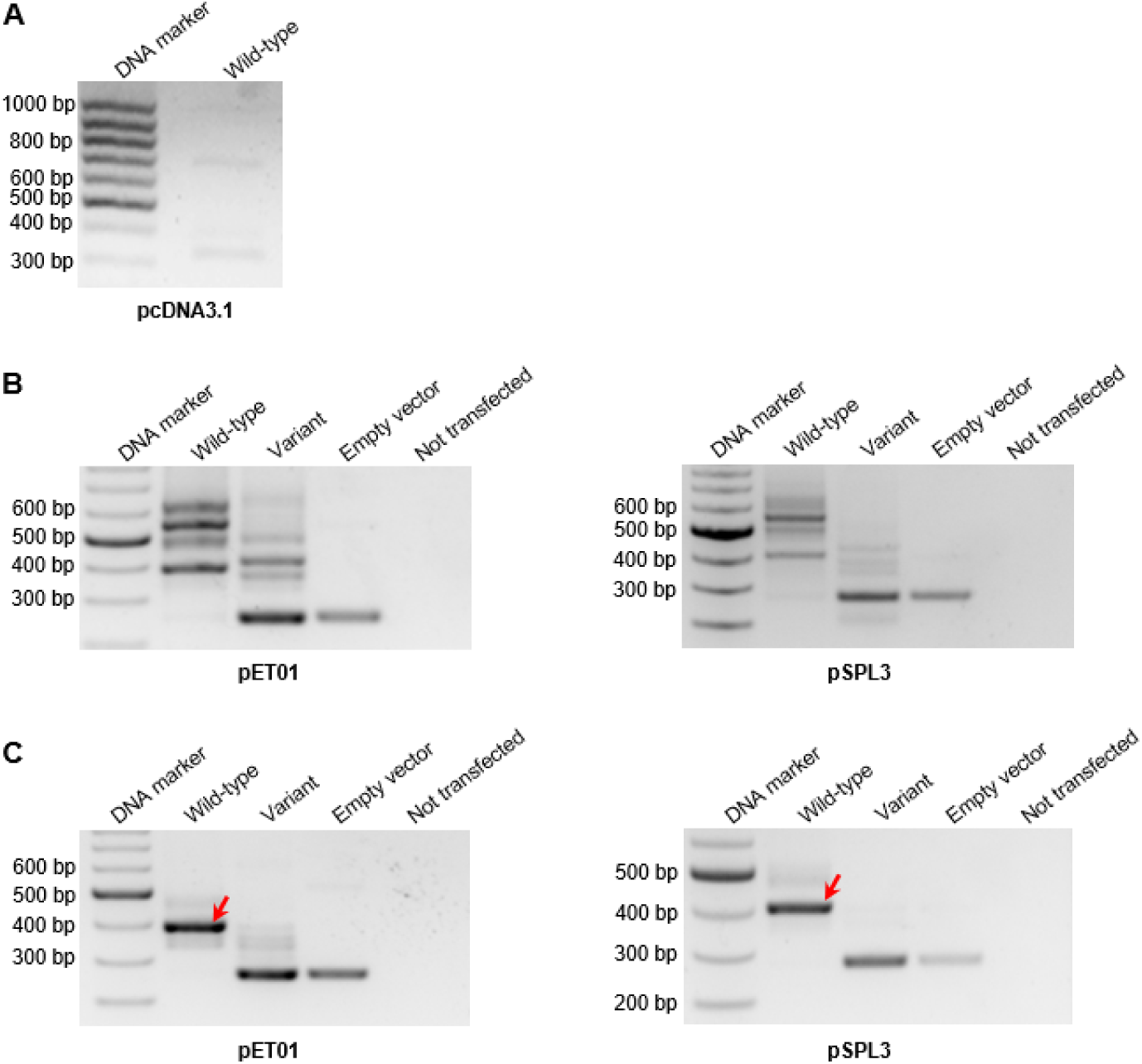
Splicing analysis of the *BAP1* c.783+2T>C variant in different assay contexts. **(A)** Full-length gene splicing assay (FLGSA). The wild-type (WT) construct failed to generate clear reverse transcription–PCR (RT–PCR) bands. **(B)** Minigene analysis of exons 8–10 (including exon 8, intron 8, exon 9, intron 9, and exon 10) with short flanking sequences from introns 7 and 10 cloned into the pET01 and pSPL3 vectors. No normally spliced transcripts were detected, in accordance with the *BAP1* mRNA reference sequence NM_004656.4, even from the WT constructs. For band annotations, see Figure S2. **(C)** Minigene analysis of exon 9 with short flanking intronic sequences cloned into the pET01 and pSPL3 vectors. Arrows indicate bands corresponding to the normally spliced transcript, in accordance with the *BAP1* mRNA reference sequence NM_004656.4. Additional bands are annotated in Figure S2. *BAP1* genomic reference sequence: NG_031859.1.

ClinVar currently lists c.783+2T>C under “conflicting classifications of pathogenicity” ([VCV000422670.29](https://www.ncbi.nlm.nih.gov/clinvar/variation/422670/); accessed October 28, 2025), with five submissions of “uncertain significance” and one of “likely benign.” In light of these inconclusive findings, allele-specific transcript analyses in heterozygous individuals—if possible in the presence of a *cis*-linked coding variant outside exon 9—would be most informative. Alternatively, targeted knock-in cell models could provide additional insight into the splicing effect of *BAP1* c.783+2T>C.

## Discussion

Our analysis yielded several insights into the impact of GT>GC 5′ splice-site variants. Consistent with findings from FLGSA,^12^ we show that disease-associated GT>GC variants can generate a wide range of residual WT transcript levels, from barely detectable to nearly normal. Moreover, our findings confirm our earlier postulate that “GT>GC variants in human disease genes may not invariably be pathogenic,”^12^ as exemplified by the *DOCK8* c.3234+2T>C variant.

Second, GT>GC variants that generate substantial yet incomplete WT transcript levels often fall within a functional grey zone, complicating their classification (Figure 5). Such indeterminacy underscores the limitations of binary frameworks and has important clinical implications. A clear example is *SPINK1* c.194+2T>C, which retains approximately 10 % of normal WT transcript. In the simple heterozygous state, its effect is too weak to be classified as “pathogenic” yet too strong to be regarded as a mere “risk” allele. However, in homozygous, compound heterozygous, or *trans*-heterozygous states—or in the presence of environmental modifiers—the same variant confers a substantially greater pathogenic effect.^8^ This exemplifies how variants with intermediate functional effects contribute to the genetic complexity of human disease. To address this challenge, we have proposed expanding the ACMG/AMP variant classification framework by incorporating additional categories, including a “predisposing” category.^6,8^

**Figure 5.**
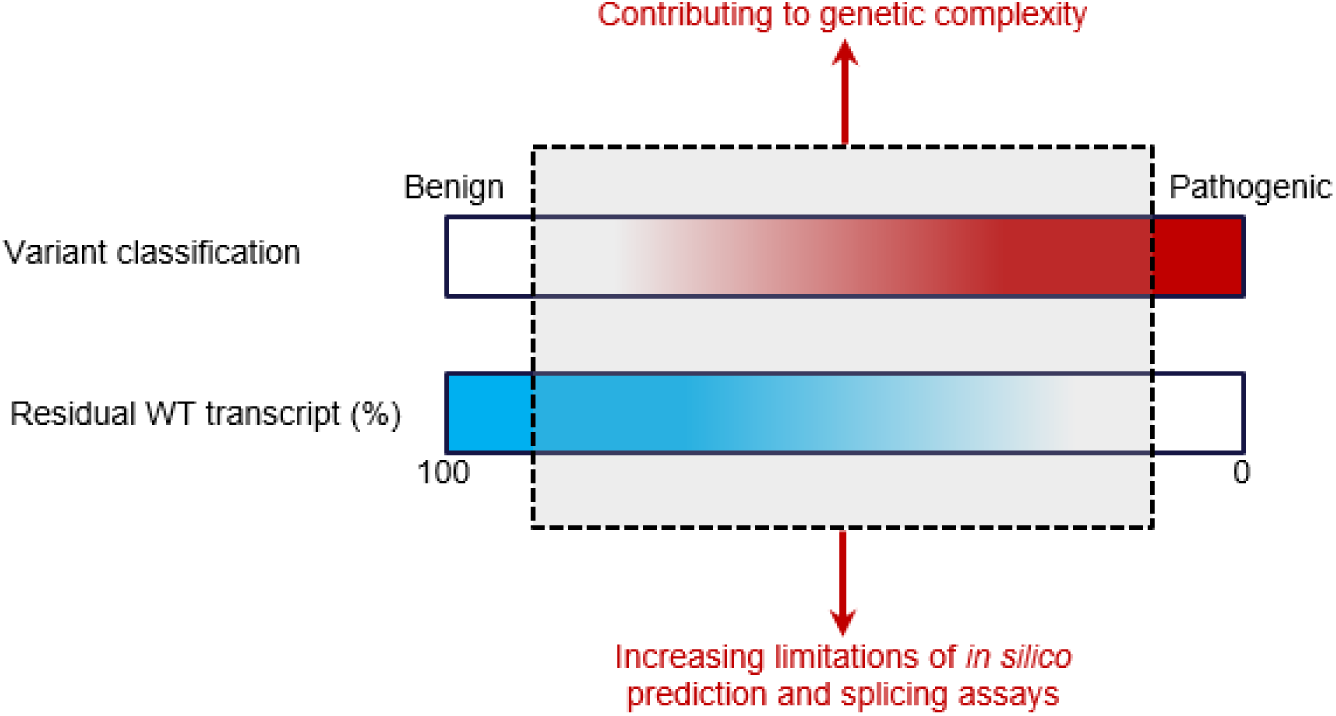
Conceptual model of the functional grey zone in variant classification. Residual wild-type (WT) transcript levels and variant classification lie along continua rather than binary states. Variants with intermediate WT transcript levels often fall within a functional grey zone, where *in silico* prediction and experimental analysis become increasingly difficult. Such grey-zone variants may act through variant–variant (both intra- and intergenic) or variant–environment interactions, thereby contributing to the genetic complexity of human disease. Although illustrated here with splice-altering variants, this conceptual model also applies to other variant types and disease mechanisms shaped by graded levels of functional impact.

Third, our findings underscore the limited resolution of current *in silico* tools, such as SpliceAI,^20^ for predicting whether GT>GC variants generate WT transcript and, if so, to what extent. Whereas most WT transcript–retaining variants exhibit low SpliceAI ΔDL scores, some have high values (>0.80), and transcript levels show no consistent correlation with predicted scores. This limitation partly reflects the intrinsic ability of GC to function as a natural donor site in approximately 1% of U2-type introns.^45^ More broadly, it highlights the difficulty of predicting splicing outcomes that are not all-or-none, leaving many variants within a functional grey zone that current algorithms cannot readily model or quantify.

Fourth, our results highlight the methodological challenges inherent in experimentally characterizing splice-altering variants. RNA from disease-relevant tissues provides the most reliable readout, yet such material is rarely available, and surrogate sources such as blood cells or lymphoblastoid lines are informative only if the gene of interest is expressed. In the absence of RNA from affected individuals, cell-based assays—FLGSA,^12^ EMG,^23^ midigene,^46^ or minigene^47^ systems—are often used, but each has intrinsic limitations. For example, not only is FLGSA impractical for large genes, but also not all successfully constructed full-length WT constructs yield the expected WT transcript. The most frequently used minigene assays are performed in a highly artificial background that lacks the natural genomic context required for proper splicing regulation.^48,49^ Such loss of context is particularly problematic for variants with intermediate effects, for which classification depends not only on detecting aberrant transcripts but also on quantifying residual WT transcript from the variant allele. Even when RNA from affected individuals is available, interpretation in heterozygotes can be confounded by WT transcript derived from the unaffected allele. For this reason, in both our previous^12^ and current studies, analyses of disease-associated GT>GC variants were focused primarily on homozygotes and hemizygotes. RNA-seq, although increasingly applied in clinical contexts,^50–52^ faces similar challenges of tissue accessibility and allele-specific quantification. Collectively, these methodological obstacles underscore the need for refined experimental systems, such as those based on transactivation or transdifferentiation.^53^

GT>GC variants that retain residual WT transcript reflect the degeneracy and context dependence of splice-site recognition. As such, findings from this study can likely be extrapolated to many other splice-altering variants—excluding those at the invariant +1, −1, and −2 positions—because *cis-*acting motifs are degenerate, and deep intronic variants can generate novel splice sites that compete with physiological ones. Although a comprehensive survey is beyond the scope of this study, several observations support this view. In SpliceVarDB, approximately 50% of variants could not be assigned unambiguously to “splice-altering” or “non–splice-altering” categories but instead fell into a “low-frequency splice-altering” group, corresponding to weak or indeterminate evidence of spliceogenicity,^54^ suggesting that grey-zone variants are common. Among 12 FLGSA-analyzed *SPINK1* coding variants showing aberrant splicing, nine (including presumed missense and synonymous variants) also generated residual WT transcript.^55^ Likewise, deep intronic variants such as *CFTR* c.1584+18672A>G (GenBank: NM_000492.4, NG_016465.4)^56^ and *CLRN1* c.254−643G>T (GenBank: NM_174878.3, NG_009168.1)^57^ have been reported to yield both aberrant and WT transcripts. Collectively, these examples indicate that residual WT transcript generation is a recurrent and underappreciated phenomenon across genes and variant types, producing outcomes along a continuum rather than at binary extremes (Figure 5). Because residual WT transcript cannot always be readily assessed in simple or compound heterozygotes, and splice-altering variants account for an estimated 10–30% of all disease-associated alleles,^11,54,58^ we infer that variants generating residual WT transcript may represent a significant source of variable penetrance and expressivity.

Nonetheless, the boundaries of the functional grey zone (Figure 5) likely vary across genes, depending on functional tolerance thresholds and modes of inheritance, such that an identical level of residual WT transcript may be clinically inconsequential in one gene–disease context yet functionally significant in another. Although developed here in the context of LoF variants for conceptual simplicity, the grey-zone model may also extend to gain-of-function or dominant-negative variants,^59^ in which disease-associated alleles within a given gene are likewise unlikely to exert uniform effects. Notably, this conceptual framework was established primarily for genes in which complete or near-complete LoF alleles cause classical Mendelian diseases. As we have previously opined,^8^ some disease-associated genes may not necessarily harbor clearly “pathogenic” variants.

Taken together, our study establishes GT>GC variants as a tractable model for addressing the classification challenges posed by splice-altering variants with intermediate functional consequences. By using residual WT transcript as a measurable parameter—and integrating clinical and genetic data, RNA analyses of material derived from affected individuals, full-length and minigene assays, and computational predictions—we show that residual WT transcript generation is a recurrent and biologically relevant phenomenon that frequently complicates classification. Gradual functional effects are also observed for other variant types, as exemplified by missense variants in *BRCA1* (MIM: 113705)^60^ and *SOD1* (MIM: 147450),^61^ both analyzed using multiplexed assays of variant effect (MAVE). These observations underscore the need to move beyond binary pathogenic/benign frameworks toward models that accommodate a continuum of effects,^6,8^ as efforts focused solely on refining ACMG/AMP criteria within a binary framework (as reflected in recent studies^62–65^) are unlikely to resolve the fundamental problem amplified by the field’s transition from a “phenotype-first” to a “genotype-first” approach.

To address this challenge, new conceptual, computational, and experimental strategies are emerging. As highlighted by Whitcomb,^66^ our recently proposed “pathogenic–predisposing–risk–benign” framework^8^ represents an important step toward resolving the variant classification challenge. The integration of machine-learning models trained on large-scale population-based datasets^9^ with functional data generated from MAVE^67,68^ holds great promise for defining the full spectrum of variant effects revealed through analyses of increasingly large cohorts.^69–71^ Ultimately, grey-zone variants may constitute a substantial portion of those currently classified as VUS or excluded for exceeding the allele-frequency thresholds set for high-penetrance disease variants. Such variants could also account for part of the “missing heritability” observed in both rare and complex diseases.^72–74^ Recognizing the widespread presence of grey-zone variants and their contribution to genetic complexity will be essential for realizing the full potential of precision medicine.^75^

Our study has several limitations. First, the number of GT>GC 5′ splice site variants known to generate substantial amounts of WT transcript remains limited. Second, the level of WT transcript generated from certain GT>GC variants, as determined by FLGSA or RT–PCR using material derived from affected individuals, may differ from that occurring under true *in vivo* conditions within the pathologically relevant tissue. Third, a potential alternative impact of GT>GC variants—namely, a reduction in overall splicing efficiency without producing aberrant transcripts—could not be adequately assessed in this analysis. Likewise, GT>GC variants that generate both WT and aberrant transcripts, but in which the aberrant transcripts are degraded by nonsense-mediated decay (NMD),^76^ could not be evaluated. These two scenarios could lead to underestimation of the true extent of partial LoF GT>GC variants. Fourth, while we highlighted the classification challenges posed by the variants discussed in detail, we did not systematically attempt to specify how each variant should be classified. This is primarily because, as we have previously noted, “defining precise thresholds between different variant categories remains challenging and must be tailored to each gene and disease context,”^8^ and because the available functional analytic data are not conclusive or fully reliable in some cases, and the associated phenotypes were not always well defined. Despite these limitations, our integrative analysis of GT>GC variants provides a coherent framework for understanding how residual WT transcript generation shapes the functional continuum of splice-altering variants and informs their broader biological and clinical interpretation.

In conclusion, viewing GT>GC variants through the lens of residual WT transcript generation offers valuable insight into how partial LoF splice-altering variants—and, more broadly, variants with intermediate functional effects—can contribute to genetic complexity, complicate both *in silico* prediction and experimental analysis, and challenge variant classification (Figure 5).

## Data and code availability

All data supporting this study are provided within the article and its supplementary materials. This study did not generate any new datasets or code.

## Acknowledgments

This study was funded by the National Natural Science Foundation of China (82000611 (to J.H.L.)), the Shanghai Pujiang Program (2020PJD061 (to J.H.L.)), and the National Key Research and Development Program of China (2025YFC2511900 (to W.B.Z). Support for this study also came from the Institut National de la Santé et de la Recherche Médicale (INSERM), France. D.N.C. wishes to acknowledge financial support from Qiagen Inc through a License Agreement with Cardiff University. The funding bodies did not play any role in the study design, collection, analysis, and interpretation of data or the writing of the article and the decision to submit it for publication.

## Author contributions

Conceptualization, J.M.C.; study design, J.H.L., H.W., W.B.Z. and J.M.C.; methodology, J.H.L., H.W., W.B.Z. and J.M.C.; experiments, J.H.L., H.W., X.Y.T. and W.B.Z.; literature search and variant collation, E.M. and J.M.C.; writing—original draft preparation, J.M.C.; writing—review and editing, J.H.L., H.W., X.Y.T., E.M., D.N.C., C.F., Z.L., W.B.Z. and J.M.C.; project administration, W.B.Z. and J.M.C.; funding acquisition, J.H.L., W.B.Z. and J.M.C. All authors have read and approved the final version of the manuscript and agree to be accountable for all aspects of the work, ensuring that questions related to the accuracy or integrity of any part of the work are appropriately investigated and resolved.

## Declaration of generative AI and AI-assisted technologies in the writing process

During the preparation of this work the authors used ChatGPT 5 in order to improve readability and language. After using this tool, the authors reviewed and edited the content as needed and take full responsibility for the content of the publication.

