## Supplementary material for "When splicing is not all or none: GT>GC 5′ splice-site variants as a model for intermediate effects and challenges in variant classification": Table S1

| Intron | c.nomenclature (NM_000492.4, NG_016465.4) | hg38 coordinate | SpliceAI ΔDL score | Variant classification or description | | | |
|---|---|---|---|---|---|---|---|
| | | | | CFTR2 | CFTR France | CFTR1 (submitted phenotype or other details as described)[a] | ClinVar |
| 1 | c.53+2T>C | 7-117480149-T-C | 0.91 | CF-causing | – | – | Pathogenic/Likely pathogenic |
| 2 | c.164+2T>C | 7-117504365-T-C | 0.87 | CF-causing | – | "The French CF patient (male) has positive sweat chloride and carries DeltaF508 on the other allele. (pers.corr. Ferec)" | Pathogenic |
| 3 | c.273+2T>C | 7-117509144-T-C | 0.86 | – | – | – | Pathogenic |
| 4 | c.489+2T>C | 7-117531116-T-C | 0.84 | CF-causing | CF-causing | "2 patients: -one carrying DelF508 on the other allele, female, born in October 1997 -one homozygous. (pers. corr. Schwarz)" | Pathogenic/Likely pathogenic |
| 5 | c.579+2T>C | 7-117534367-T-C | 0.95 | CF-causing | – | – | – |
| 6 | c.743+2T>C | 7-117535413-T-C | 0.95 | CF-causing | – | "nasal polyps no GI symptoms no history of chest infection till now but on off cough 2 brothers with same mutation one is 10 years old with sweat chloride test is 94 and 91, on off cough no pneumonia, nasal polyps and sputum culture is staphylococcus no GI problems. other child is 3 years old with sweat chloride test 85 no pneumonia no GI problems" | Likely pathogenic |
| 7 | c.869+2T>C | 7-117536675-T-C | 0.98 | – | – | – | – |
| 8 | c.1116+2T>C | 7-117540348-T-C | 0.88 | CF-causing | – | – | Pathogenic/Likely pathogenic |
| 9 | c.1209+2T>C | 7-117542110-T-C | 0.97 | – | – | – | – |
| 10 | c.1392+2T>C | 7-117548825-T-C | 0.98 | – | – | – | – |
| 11 | c.1584+2T>C | 7-117559657-T-C | 0.84 | CF-causing | CF-causing | "The CF female patient died at 16 years of age. She was PI, with severe lung disease and carried deltaF508 on the other allele. (pers.corr. Claustres)" | Pathogenic/Likely pathogenic |
| 12 | c.1679+2T>C | 7-117587835-T-C | 0.98 | CF-causing | – | "Late diagnosis of CF in a ten year-old boy; positive sweat tests; recurrent chest infections; clubbing; history of abdominal pain and constipation" | – |
| 13 | c.1766+2T>C | 7-117590441-T-C | 0.86 | CF-causing | – | "Steatorrhea" | Pathogenic |
| 14 | c.2490+2T>C | 7-117592659-T-C | 0.35 | – | VUS (non-CF; reported in two asymptomatic homozygotes) | "Healthy individual (this mutation was identified at the heterozygous state in the partner of a CBAVD patient. The woman is Algerian)" | Conflicting classifications of pathogenicity (Pathogenic × 3; Likely pathogenic × 2; Uncertain significance × 1) |
| 15 | c.2619+2T>C | 7-117595060-T-C | 0.89 | CF-causing | – | – | Pathogenic |
| 16 | c.2657+2T>C | 7-117602865-T-C | 0.85 | – | – | – | – |

| | | | | | | | |
|---|---|---|---|---|---|---|---|
| 17 | c.2908+2T>C | 7-117603784-T-C | 0.86 | – | – | "Mutation 3040+2T->C, was identified in a CF patient who carries the [delta]F508 mutation on the other chromosome; this mutation alters the second nucleotide of intron 15 (splicing donor site). The above mutation was identified by DGGE analysis and confirmed by DNA sequencing."<br>(N.B. Classified by us as "pathogenic," as it occurs in *trans* with another "pathogenic" variant and was identified in a CF patient) | – |
| 18 | c.2988+2T>C | 7-117606755-T-C | 0.99 | – | – | "A 12 years old Chinese girl presented chronic productive cough with bilateral bronchiectasis and pancreatic insufficiency. Sweat chloride was 108 mmol/L. The ultrasound of abdomen showed liver cirrhosis and splenomegaly. The patient was also heterozygous for W679X mutation. Her patients had normal phenotype."<br>(N.B. Classified by us as "pathogenic," as it occurs in *trans* with another "pathogenic" variant and was identified in a CF patient) | Pathogenic |
| 19 | c.3139+2T>C | 7-117610671-T-C | 0.99 | CF-causing | – | – | – |
| 20 | c.3367+2T>C | 7-117611810-T-C | 0.98 | CF-causing | – | "A mutation at 3499+2 T->C which may affect the splice site, in a CF patient who carries a [delta]F508 on the other chromosome. The mutation has been confirmed by dot blots with a specific oligonucleotide." | – |
| 21 | c.3468+2T>C | 7-117614715-T-C | 0.98 | – | Likely pathogenic | "This sequence variation was detected by DGGE and identified by direct sequencing. This splice mutation 3600+2T->C was not found in 100 other non-[delta]F508 CF chromosomes and 500 non-CF chromosomes tested. This mutation was observed in a male subject with congenital bilateral absence of the vas deferens. The chloride sweat test was normal (40 mM), but the patient had abnormal nasal potential difference. The other mutation is unknown." | Pathogenic |
| 22 | c.3717+2T>C | 7-117627772-T-C | 0.56 | – | – | – | – |
| 23 | c.3873+2T>C | 7-117642595-T-C | 0.98 | CF-causing | CF-causing | "4005+2T->C is one of the most commonly encountered in Norwegian CF patients; this mutation had not been seen in Denmark or in Sweden. (pers. corr. Boman)" | Pathogenic |
| 24 | c.3963+2T>C | 7-117652933-T-C | 0.98 | – | – | – | – |
| 25 | c.4136+2T>C | 7-117664862-T-C | 0.97 | – | – | – | – |
| 26 | c.4242+2T>C | 7-117665566-T-C | 0.15 | – | – | "2 patients with suspected diagnosis of CF" | Pathogenic/Likely pathogenic |

**Table S1. All possible GT>GC variants in *CFTR***

[a]In contrast to CFTR2 and CFTR France, variants in CFTR1 are not classified according to pathogenicity.

The three variants with low SpliceAI ΔDL scores are shaded in yellow. For the two variants classified by this study as "pathogenic," the corresponding reasons are provided in parentheses after the clinical and genetic descriptions (highlighted in red) of the affected individuals in CFTR1.

Abbreviations: CBAVD, congenital bilateral absence of the vas deferens; CF, cystic fibrosis; ΔDL, delta donor loss; DGGE, denaturing gradient gel electrophoresis; GI, gastrointestinal; PI, pancreatic insufficiency; VUS, variants of uncertain significance.
