## Supplementary material for "When splicing is not all or none: GT>GC 5′ splice-site variants as a model for intermediate effects and challenges in variant classification": Figure S1

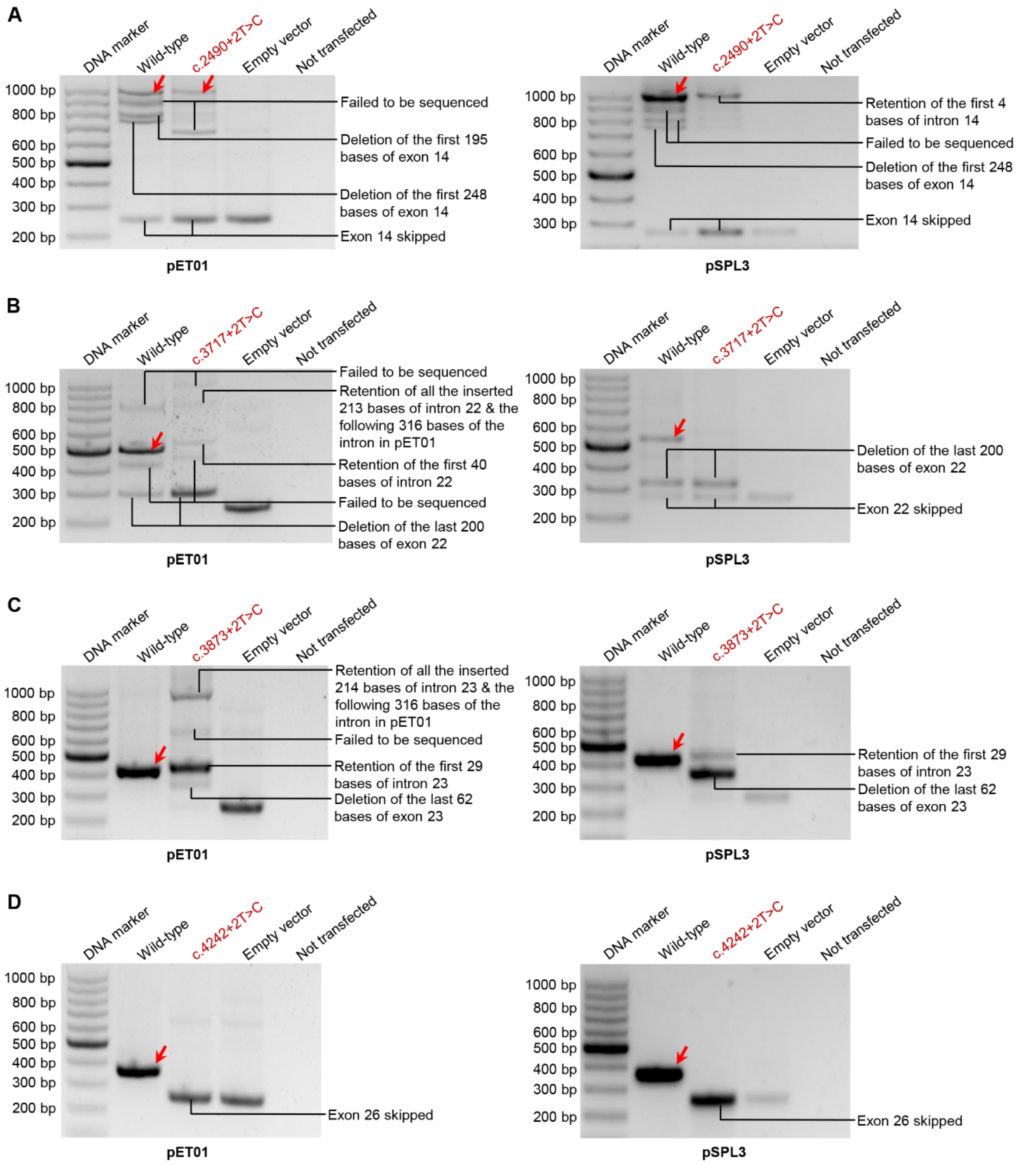


**Figure S1. Minigene splicing analysis of four selected *CFTR* GT>GC substitutions using pET01 and pSPL3 vectors**

This figure expands on Figure 3 and provides full annotation of all observed bands.

Each variant was analyzed with its preceding exon and short flanking intronic sequences cloned into the respective reporter vectors. Arrows indicate bands corresponding to the normally spliced transcript, in accordance with the *CFTR* mRNA reference sequence NM_000492.4.
