## Supplementary material for "When splicing is not all or none: GT>GC 5′ splice-site variants as a model for intermediate effects and challenges in variant classification": Figure S2

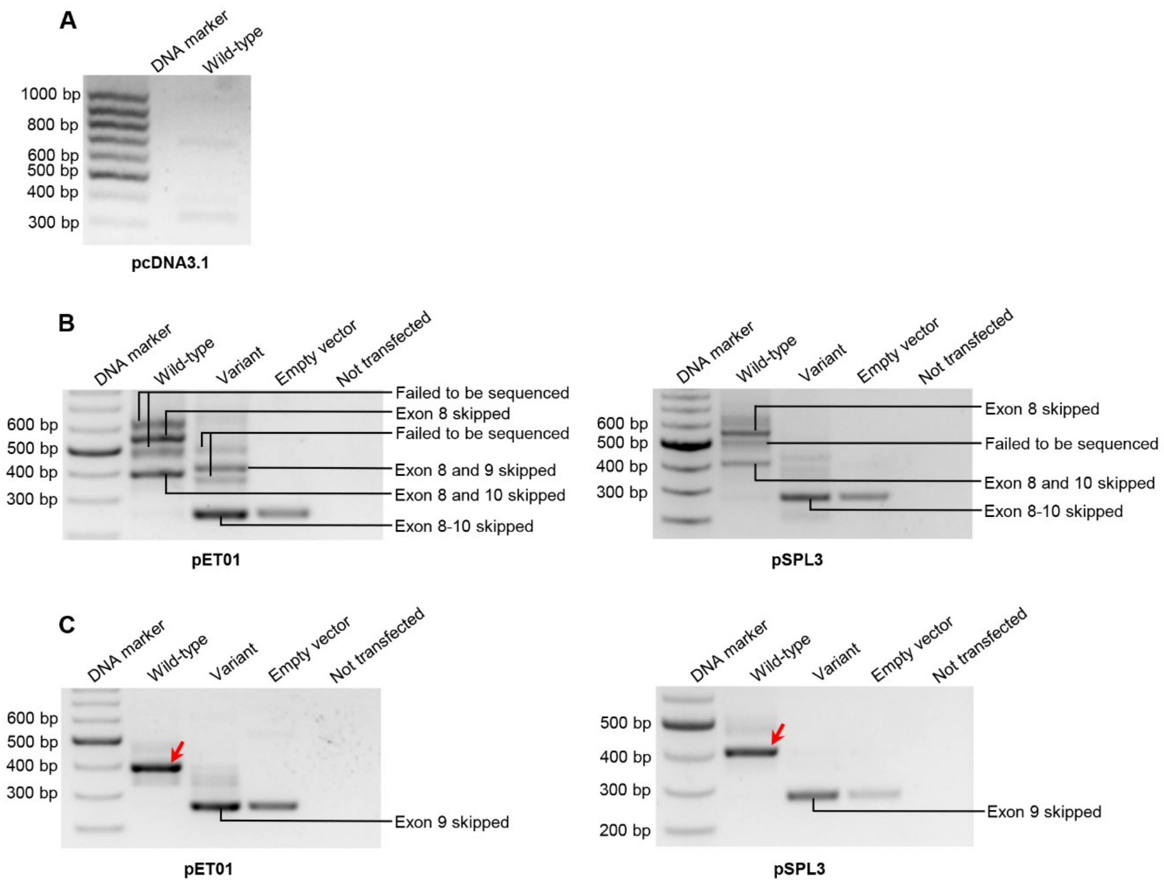


**Figure S2. Splicing analysis of the *BAP1* c.783+2T>C variant in different assay contexts**

This figure expands on Figure 4 and provides full annotation of all observed bands.

(**A**) Full-length gene splicing assay (FLGSA). The wild-type (WT) construct failed to generate clear reverse transcription–PCR (RT–PCR) bands.

(**B**) Minigene analysis of exons 8–10 (including exon 8, intron 8, exon 9, intron 9, and exon 10) with short flanking sequences from introns 7 and 10 cloned into the pET01 and pSPL3 vectors. No normally spliced transcripts were detected, in accordance with the *BAP1* mRNA reference sequence NM_004656.4, even from the WT constructs.

(**C**) Minigene analysis of exon 9 with short flanking intronic sequences cloned into the pET01 and pSPL3 vectors. Arrows indicate bands corresponding to the normally spliced transcript, in accordance with the *BAP1* mRNA reference sequence NM_004656.4.

*BAP1* genomic reference sequence: NG_031859.1.
